# Innate Immune Priming by cGAS as a Preparatory Countermeasure Against RNA Virus Infection

**DOI:** 10.1101/434027

**Authors:** Michael T. Parker, Smita Gopinath, Corey E. Perez, Melissa M. Linehan, Jason M. Crawford, Akiko Iwasaki, Brett D. Lindenbach

## Abstract

The detection of nucleic acids by pattern recognition receptors is an ancient and conserved component of the innate immune system. Notably, RNA virus genomes are sensed by mammalian cytosolic RIG-I–like receptors, thereby activating interferon-stimulated gene (ISG) expression to restrict viral replication. However, recent evidence indicates that the cGAS-STING DNA sensing pathway also protects against RNA viruses. So far, the mechanisms responsible for DNA sensing of RNA viruses, which replicate without known DNA intermediates, remain unclear. By using cGAS gene knockout and reconstitution in human and mouse cell cultures, we discovered that DNA sensing and cGAMP synthase activities are required for cGAS-mediated restriction of vesicular stomatitis virus and Sindbis virus. The level of cGAMP produced in response to RNA virus infection was below the threshold of detection, suggesting that only transient and/or low levels of cGAMP are produced during RNA virus infections. To clarify the DNA ligands that activate cGAS activity, we confirmed that cGAS binds mitochondrial DNA in the cytosol of both uninfected and infected cells; however, the amount of cGAS-associated mitochondrial DNA did not change in response to virus infection. Rather, a variety of pre-existing cytosolic DNAs, including mitochondrial DNA and endogenous cDNAs, may serve as stimuli for basal cGAS activation. Importantly, cGAS knockout and reconstitution experiments demonstrated that cGAS drives low-level ISG expression at steady state. We propose that cGAS-STING restricts RNA viruses by promoting a preparatory immune activation state within cells, likely primed by endogenous cellular DNA ligands.

**Importance:** Many medically important RNA viruses are restricted by the cGAS-STING DNA-sensing pathway of innate immune activation. Since these viruses replicate without DNA intermediates, it is unclear what DNA ligand(s) are responsible for triggering this pathway. We show here that cGAS’s DNA binding and signaling activities are required for RNA virus restriction, similar to the mechanisms by which it restricts DNA viruses. Furthermore, we confirmed that cGAS continuously binds host DNA, which was unaffected by RNA virus infection. Finally, cGAS expression correlated with the low-level expression of interferon-stimulated genes in uninfected cells, both *in vitro* and *in vivo*. We propose that cGAS-mediated sensing of endogenous DNA ligands contributes to RNA virus restriction by establishing a baseline of innate immune activation.

## Introduction

A key feature of innate immunity is the detection of pathogen-associated molecular patterns (PAMPs) by pattern recognition receptors (PRRs) (1). For mammalian cells, viral nucleic acids are detected by distinct PRRs, triggering interferon-stimulated gene (ISG) expression to set up an antiviral state. During RNA virus infections, uncapped and double-stranded RNAs are detected in the cytosol by the PRRs retinoic acid-inducible gene I (RIG-I) and related RIG-I-like receptors (RLRs). However, the recent discovery of the cGAS-STING cytosolic DNA sensing pathway, and the observation that it can also restrict RNA viruses (2), reveals a need to further investigate the mechanisms of nucleic acid sensing during RNA virus infection.

The stimulator of interferon genes (STING) is an endoplasmic reticulum- and mitochondrial-bound protein that spontaneously activates ISG expression when overexpressed (2). Although STING is involved in DNA sensing, STING^−/−^ mice and mouse endothelial fibroblasts (MEFs) are more permissive for vesicular stomatitis virus (VSV), a negative-stand RNA virus (2, 3). Additionally, studies in MEFs deficient in three prime repair exonuclease 1 (TREX1), a nuclease important for the turnover of cytosolic retroelement cDNAs (4), have described enhanced antiviral phenotypes in response to a wide array of RNA viruses and retroviruses, presumably due to the accumulation of DNA in the cytosol (5, 6). It appears that this DNA-based restriction is broad, as many RNA viruses have evolved mechanisms to subvert the cGAS-STING pathway, including flaviviruses (7–9), hepaciviruses (10, 11), picornaviruses (3), coronaviruses (12–17), and influenza A virus (18).

STING does not directly interact with cytosolic DNA, but functions as an innate immune adaptor protein to transduce signals between cyclic GMP-AMP synthase (cGAS) and Tank-binding kinase 1, which subsequently phosphorylates the transcription factor interferon regulatory factor 3 (IRF3) to initiate an ISG response (19). Recent evidence also suggests that STING inhibits translation by unknown mechanisms and may restrict RNA virus replication independent of IRF3 activation (20).

cGAS is a nucleic acid-binding protein specific for dsDNA and DNA:RNA hybrids that also has nucleotidyl transferase activity (21–24). DNA binding induces structural changes to form the cGAS active site, which synthesizes a non-canonical 5′–2’- and 5′–3’-linked cyclic dinucleotide known as cyclic guanosine monophosphate– adenosine monophosphate (cGAMP) (25–28). cGAMP is a diffusible secondary messenger that specifically binds to STING with high affinity (*K_D_* ~4 nM), thereby inducing a downstream innate immune response (29–32).

For RNA viruses that replicate in the cytosol without a DNA intermediate, the specific ligands that activate cGAS remain unclear. At present, the prevailing hypothesis is that RNA viruses induce release of mitochondrial DNA (mtDNA) into the cytosol, thereby activating innate immune responses (7, 33–36). However, it is unclear whether mitochondrial damage is a conserved feature of RNA virus infection, nor is it clear that cGAS-STING activation follows the same pathway for both RNA and DNA viruses.

In this study, we investigated whether the DNA binding and cGAMP synthesis activities of human cGAS (hcGAS) are required for RNA virus restriction. While both activities were required, the amount of cGAMP produced during virus infection was too low to detect. We also confirmed that hcGAS binds mtDNA in both uninfected and infected cells but did not observe increased cytosolic or cGAS-associated mtDNA in response to RNA virus infection. We found that cGAS stimulated smoldering, low-level innate immune activation, most likely in response to endogenous DNA ligands, suggesting that cGAS-STING can passively restrict incoming RNA viruses.

## Results

### cGAS mediates restriction of RNA viruses in immortalized MEFs

To clarify the role of cGAS in restriction of RNA virus replication, we performed viral single-step growth curve experiments in wild-type (WT) and cGAS^−/−^ (KO) MEFs immortalized with SV40 large T antigen (Figure 1). Both VSV-GFP and SINV-GFP grew to higher titers in KO MEFs (Figure 1A, B). We then asked whether reconstituting cGAS expression in KO MEFs could restore RNA virus restriction by performing VSV plaque assays on WT MEFs, KO MEFs, or KO MEFs stably expressing hcGAS-HA3x, a functional, triple HA-tagged form of human cGAS (34). As seen in Figure 1C, both WT and hcGAS-reconstituted (KO+WT) cells significantly reduced VSV-GFP plaque formation compared to KO MEFs. These results confirm previous observations that cGAS can restrict RNA virus infection.

**Figure 1.**
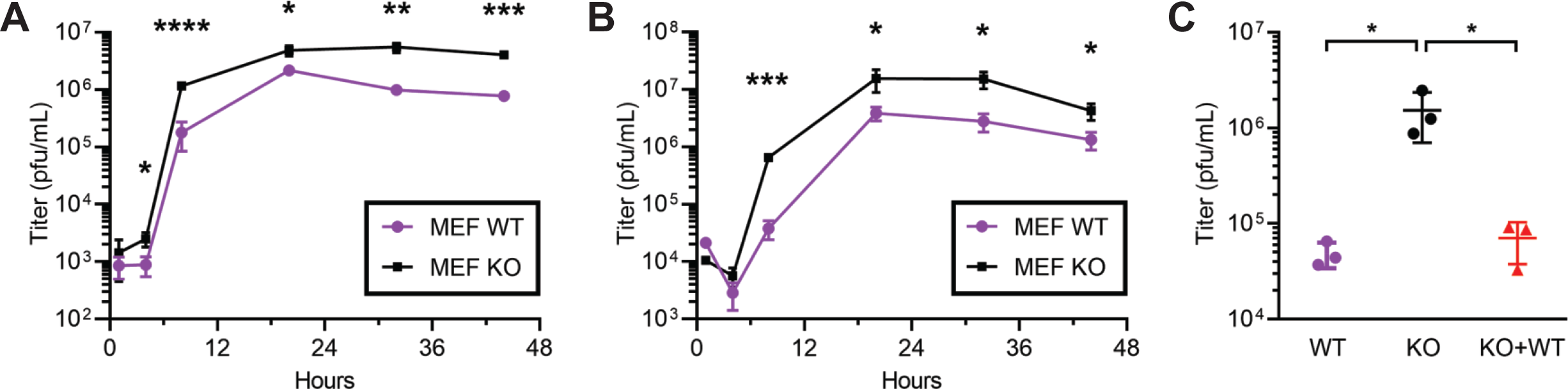
WT cGAS and hcGAS restrict RNA virus infection in immortalized MEFs. Single-step growth curve of (A) VSV-GFP and (B) SINV-GFP production from MEFs infected at MOI 3, as assessed by plaque assay on SW13 cells. (C) VSV-GFP stocks were titered by plaque assay on WT, KO, and KO+WT cells.

### The cGAS DNA binding- and cGAMP synthase active site residues are essential for RNA virus restriction

It is currently unclear whether cGAS restricts RNA viruses via the same mechanism that it restricts DNA viruses. We therefore asked whether the DNA binding and cGAMP synthase activities, which are required for DNA sensing and downstream STING activation, are also required for cGAS-mediated restriction of RNA viruses. Specifically, we reconstructed previously described loss-of-function mutations in the DNA binding pocket and cGAMP synthase active site within hcGAS-HA3x (27) (Figures 2A and S1), then restored cGAS expression in KO MEFs, as above. Notably, expression levels of hCGAS-HA3x were similar to endogenous mouse cGAS (Figure 2B). As expected, WT hcGAS-HA3x expression reduced VSV-GFP production, while expression of the DNA binding and catalytically inactive hcGAS-HA3x mutants did not (Figure 2B). These results indicate that cGAS-mediated restriction of an RNA virus depends on its DNA binding and cGAMP synthase activities.

**Figure 2.**
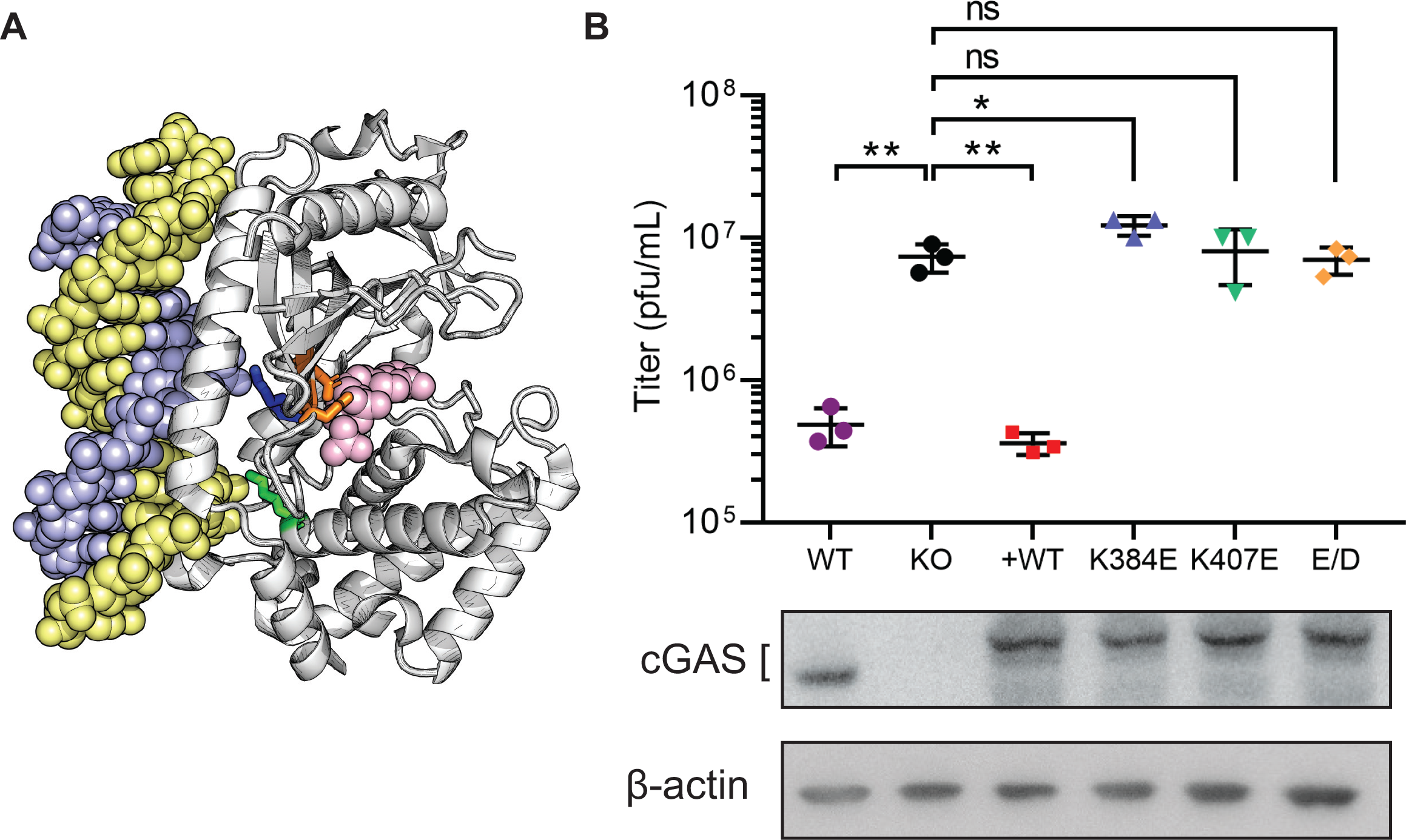
hcGAS-HA3x variants display differential restriction of RNA virus infection. (A) The structure of hcGAS (grey) in complex with ATP (pink) and DNA (yellow-purple), as rendered from PDB 6CTA; residues K384 (blue), K407 (green), and E225/D227 (orange) are shown. (B) Plaque assay of VSV-GFP produced by SV40 T-immortalized MEF KO cells reconstituted with hcGAS-HA3x variants and infected (MOI 3) for 8 hours; western blot of hcGAS expression is shown below.

Because SV40 T antigen and other viral oncogenes can inhibit innate immune responses, including cGAS-STING activation (37), we sought to confirm the above findings in untransformed cells. We therefore reconstituted primary cGAS^−/−^ MEFs with WT or mutant forms of hcGAS-HA3x and then assessed their ability to restrict the growth of VSV-GFP, SINV-GFP, or VSVΔM51A-GFP, a VSV mutant (M51A in the M gene) that is more susceptible to innate immune responses (38). All three viruses were significantly restricted in primary MEFs reconstituted with WT hcGAS-HA3x but not with the DNA-binding nor cGAMP-synthase active site mutants (Figure 3A-C). Restriction of VSVΔM51A-GFP was more potent than VSV-GFP. SINV-GFP was potently restricted by hcGAS WT but not by either DNA binding mutant; SINV-GFP infection was modestly reduced in cells expressing the E225A/D227A mutant.

**Figure 3.**
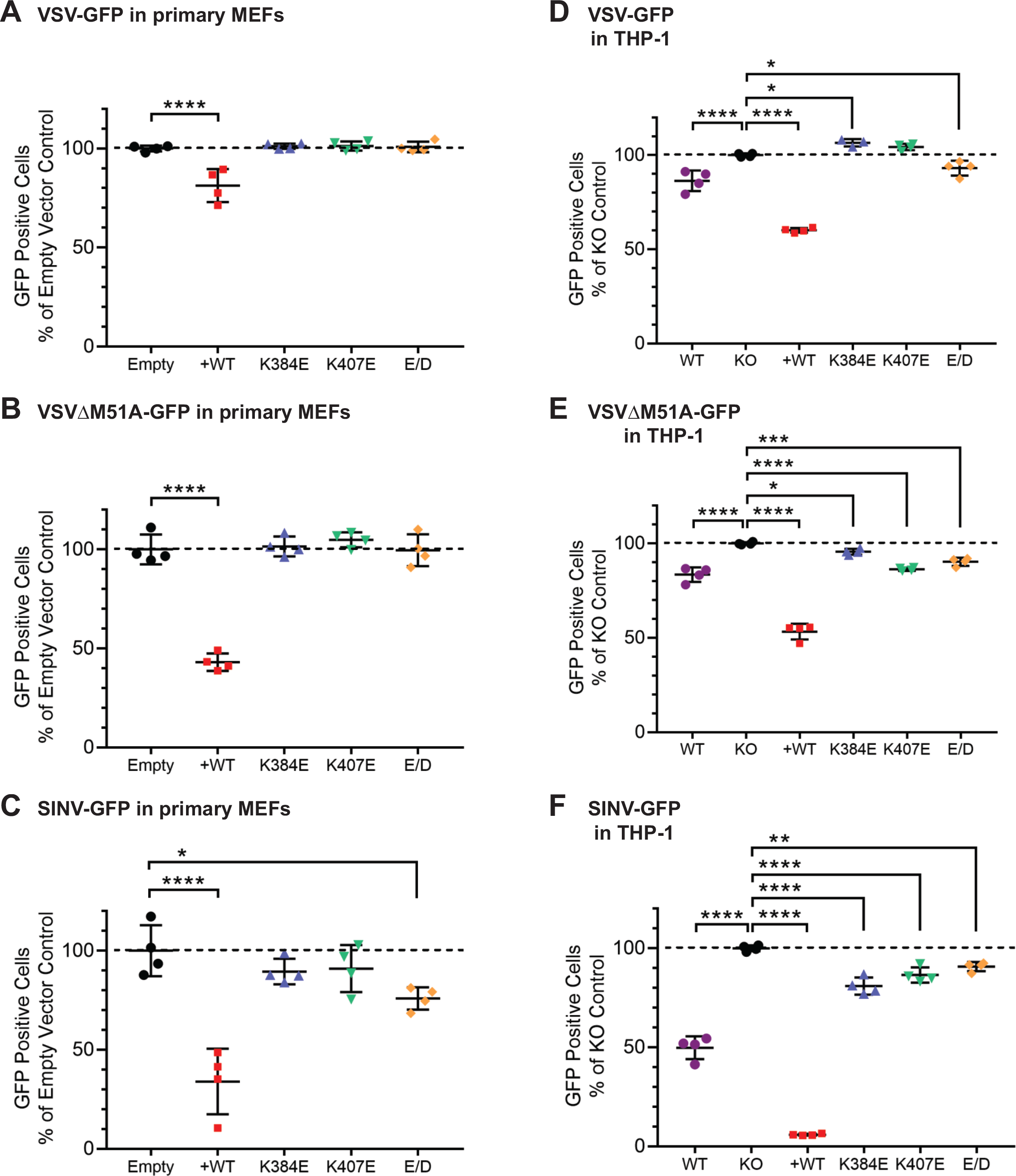
hcGAS variants display differential restriction of RNA virus infection. Primary KO MEFs were transduced to express hcGAS variants with lentiviral vectors and infected with (A) VSV-GFP, (B) VSVΔM51A-GFP, or (C) SINV-GFP; % infected cells was determined by flow cytometry in relation to empty vector-transduced KO MEFs (Empty). WT THP-1, THP-1 cGAS knockout (KO), and THP-1 cGAS KO cells reconstituted with WT (+WT) mutant forms of hcGAS (K384E, K407E, or E/D) were differentiated with PMA and infected with (D) VSV-GFP, (E) VSVΔM51A-GFP, or (F) SINV-GFP; % infected cells was determined by flow cytometry in relation to THP-1 KO cells.

To further corroborate the role of cGAS in restriction of RNA viruses in immunocompetent human cells, we utilized the THP-1 human monocyte line that has robust DNA sensing capability (21). First, we used CRISPR/Cas9 to generate cGAS KO THP-1 monocytes, then established stable lines reconstituted with WT or mutant hcGAS-HA3x; it should be noted that hcGAS-HA3x was overexpressed 2- to 6-fold in THP-1 cells relative to endogenous hcGAS (Figure S2). Differentiated WT THP-1 cells and THP-1 KO cells reconstituted with WT hcGAS-HA3x restricted growth of VSV-GFP, VSVΔM51A-GFP, and SINV-GFP, while THP-1 KO cells or THP-1 KO cells reconstituted with inactive hcGAS-HA3x mutants showed little or no restriction (Figure 3D–3F). As observed previously in MEFs, VSVΔM51A-GFP was more potently restricted than VSV-GFP, but unlike in MEFs, infected fewer cells expressing mutant cGAS. This was also true for SINV-GFP, albeit restriction with WT hcGAS-HA3x was extremely potent, comparatively. It is unclear whether these modest decreases in infection of the cGAS mutants was due to hcGAS-HA3x overexpression in THP-1 cells, residual hcGAS activities, or normal clonal variation of cells. Nevertheless, these results are most consistent with an integral role for cGAS DNA binding and cGAMP synthase activities in RNA virus restriction.

### Detection of cGAMP produced in response to DNA transfection but not RNA virus infection

Because cGAMP synthesis activity was essential for RNA virus restriction, we next sought to identify cGAMP produced in response to RNA virus infection or, as a positive control, DNA transfection, by using liquid chromatography-mass spectrophotometry (LC-MS) and LC-MS/MS. HEK 293E cells were used in these experiments because this cell line lacks endogenous cGAS expression and could be reconstituted with WT or mutant hcGAS-HA3x; however, unlike MEFs and THP-1 KO cells, HEK 293E cells could be efficiently transfected with DNA and readily scaled up for isolation of cGAMP from cytosolic extracts. As shown in Figure 4A, a unique UHPLC peak (~5 minutes elution) was observed after transfecting WT hcGAS-HA3x-expressing HEK 293E cells with salmon sperm DNA; MS analysis confirmed that this peak corresponded to cGAMP (Figures 4B and 4C). Moreover, cGAMP was not observed in untransfected cells expressing WT cGAS or in DNA-transfected cells expressing a catalytically inactive form of hcGAS (Figure 4D). Surprisingly, cGAMP remained below detectable levels after 5 hours of VSV-GFP infection at a MOI of 10 (Figure 4D), suggesting that detectable levels of cGAMP were not produced in response to RNA virus infection.

**Figure 4.**
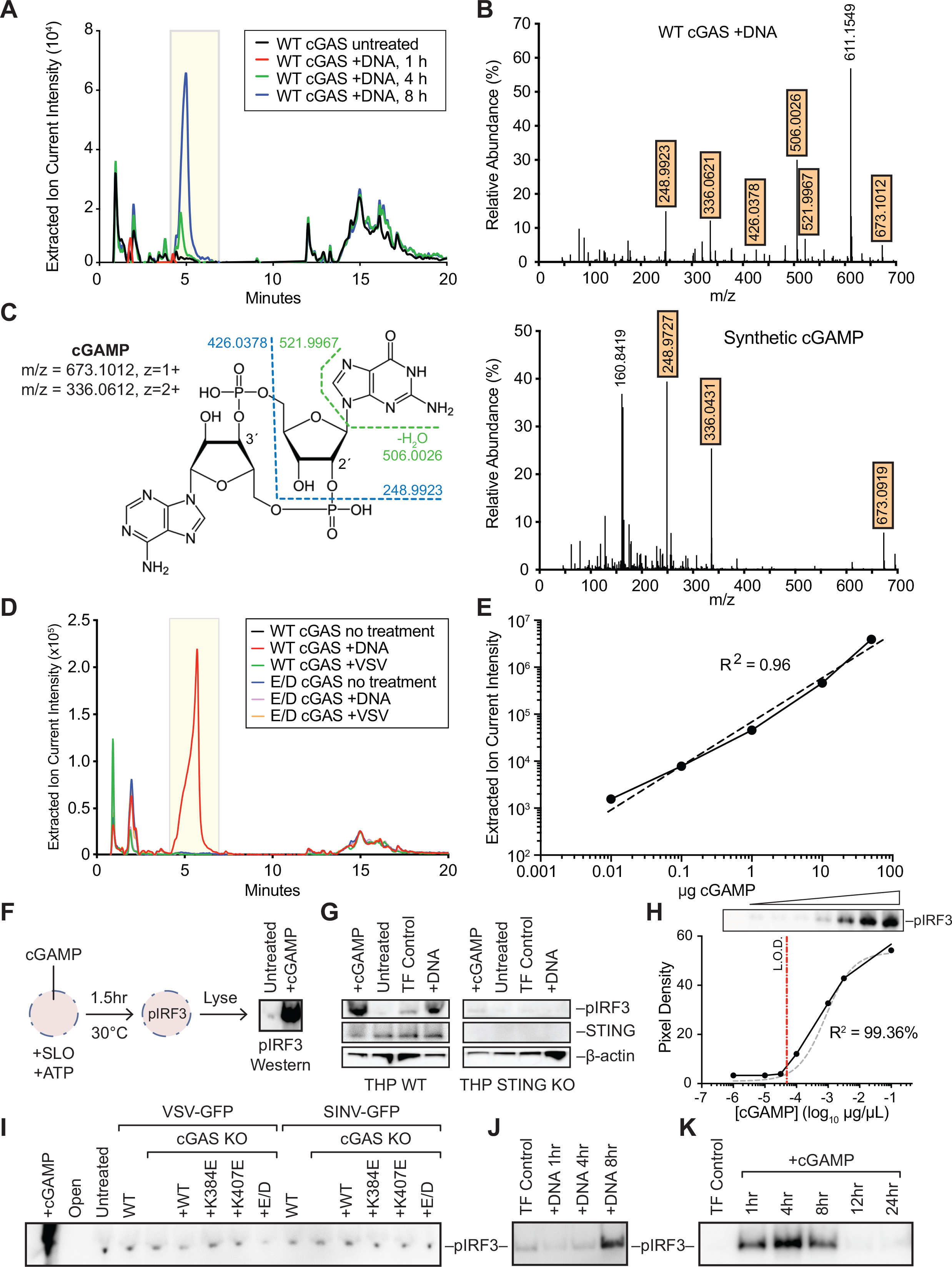
VSV-GFP infection does not induce detectable cGAMP production. (A) UHPLC profiles showing a time-course of cGAMP production after transfecting salmon sperm DNA into hcGAS-3xHA–expressing HEK 293E cells. The yellow box represents the peak elution range of synthetic cGAMP observed in pilot experiments. (B) Mass chromatogram of the eluted cGAMP peak after transfecting DNA hcGAS-3xHA–expressing HEK 293E cells. Known ionization products of cGAMP are highlighted in orange. (C) Diagram of cGAMP indicating predicted fragmentation pattern from MS data of cell-derived cGAMP; for reference a mass chromatogram obtained from synthetic cGAMP (Invivogen) is shown. (D) UHPLC profiles of untreated, VSV infected, or DNA transfected HEK293E cells expressing WT or catalytically inactive hcGAMP-3xHA. (E) Standard curve of extracted ion currents vs. synthetic cGAMP input. (F) Workflow of the cGAMP bioassay, see text for details. (G) cGAMP-mediated IRF3 phosphorylation is dependent on STING. WT or STING KO THP-1 were transfected with cGAMP, DNA, or left untransfected; TF Controls received transfection reagent but no DNA. pIRF3, STING, and ß-actin were detected by western blot. (H) Standard curve of pIRF3 detection vs. synthetic cGAMP input; L.O.D., limit of detection. (I) cGAMP was not detected during RNA virus infections. WT THP-1 or cGAS KO cells expressing the indicated forms of hcGAS-HA3x were infected with VSV-GFP or SINV-GFP at MOI 3 for 5 hours. Data are representative of multiple experiments performed at various scales and lengths of infection. (J) Time-course of cGAMP formation after transfecting DNA into HEK 293E cells expressing hcGAS-HA3x. (K) Time-course of cGAMP activity in whole cell lysates of HEK 293E cells transfected with cGAMP.

While the LC/MS technique provides exquisite specificity for identifying cGAMP in complex cytosolic extracts, cGAMP biological assays may be more sensitive. Indeed, our UPLC-MS configuration reliably detected nanogram amounts of cGAMP spiked into cytosolic extract (Fig. 4E), which equates to >1 million molecules of cGAMP per cell. We therefore established a bioassay for cGAMP-mediated IRF-3 activation in streptolysin O-(SLO)-permeabilized cells (Figure 4F). This bioassay was shown to be dependent on STING activation (Fig. 4G) and had a limit of detection (L.O.D.) of ~5 x 10^−4^ μg/μl (~0.74 μM) cGAMP (Figure 4H), in line with other published cGAMP bioassays (21). Again, we were unable to detect cGAMP in lysates from VSV-infected or SINV-infected THP-1 cells expressing WT hcGAS, while a synthetic cGAMP control led to robust phosphorylation of IRF3 (Figure 4I). To validate that cell-derived cGAMP could be detected by this assay, a time-course experiment was conducted by transfecting HEK 293E cells expressing WT hcGAS with salmon sperm DNA, revealing the time-dependent increase in cGAMP (Figure 4J). Furthermore, we found that transfected cGAMP was rapidly turned over within hours (Fig. 4K), most likely via the ENPP1 phosphodiesterase previously reported to turnover cGAMP in mammalian cells (39). Collectively, these results indicate that if cGAMP is produced in response to RNA virus infection, it may be produced at levels below the limit of our detection and/or rapidly turned over.

### cGAS binds mitochondrial DNA at steady state and during RNA virus infection

Given that cGAS DNA binding activity was also required for RNA virus restriction, we sought to identify DNA ligands of cGAS during RNA virus infection. First, we identified conditions to specifically co-immunoprecipitate cGAS and mtDNA, a known DNA ligand (34). As shown in Figure 5A, mtDNA was specifically enriched by HA-immunoprecipitation from cells expressing WT hcGAS-HA3x, but not from cells expressing the K384E DNA binding mutant. It should be noted that this experiment is representative of many iterations performed at different scales. Given prior links between virus infection, mitochondrial stress, and cGAS-mtDNA interaction (34, 40), we next asked whether VSV altered the amount of cGAS-associated mtDNA. Surprisingly, VSV-GFP infection had no impact on the amount of cGAS-associated mtDNA (Fig. 5B), which led us to isolate cytosolic DNA (Figure 5C) to quantitate mtDNA content with and without infection. Unexpectedly, VSV-GFP infection had no impact on either the total amount of cellular mtDNA (Fig. 5D) or cytosolic mtDNA (Fig. 5E).

**Figure 5.**
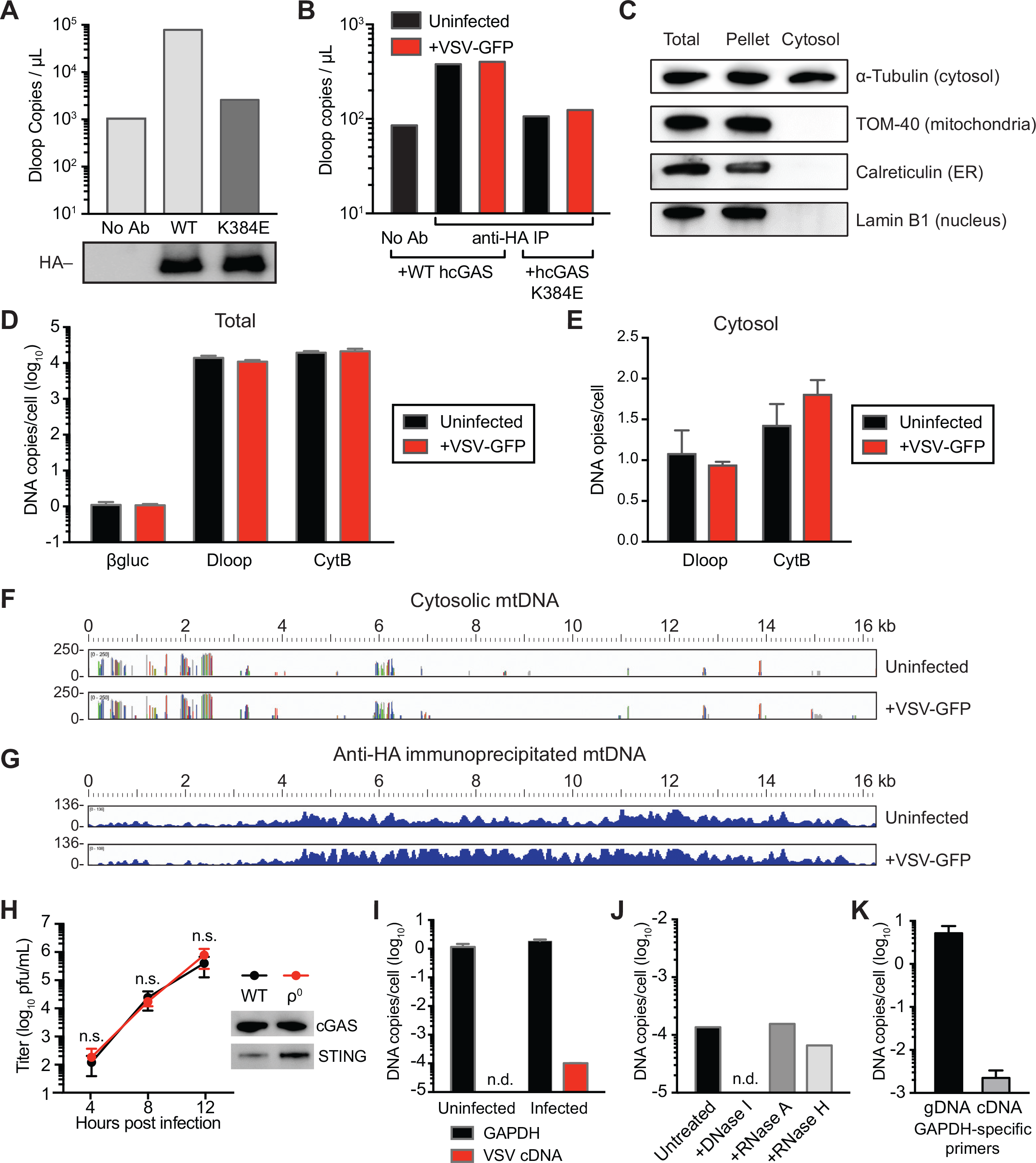
VSV infection does not introduce cGAS DNA ligands. (A) Isolation of cGAS-bound mtDNA. The amount of mtDNA D-loop sequence was quantitated by qPCR after HA-immunoprecipitation from MEF cGAS KO cells reconstituted with WT hcGAS-HA3x or the K384E DNA binding mutant. The No Ab control was from WT cells. This experiment was repeated many times at different scales, with similar cGAS-specific enrichment of mtDNA. (B) The mtDNA content of VSV-GFP-infected and uninfected MEFs was assessed by D-loop qPCR. (C) Western blotting of organelle/compartment-specific proteins in MEF WT total and cytosolic fractions with 25μg/mL digitonin extraction. (D) Total amounts of mtDNA (Dloop and CytB) and cellular DNA (ß-gluc) were determined by qPCR in uninfected and VSV-GFP-infected MEF cells. (E) The mtDNA content was determined in cytosolic extracts from uninfected and VSV-GFP-infected MEF cells. (F) Deep sequencing of cytosolic extracts from uninfected and VSV-GFP-infected MEFs revealed the presence of mtDNA. (G) Deep sequencing of cGAS-immunoprecipitates from uninfected and VSV-GFP-infected reveal abundant mtDNA. (H) Time course of VSV-GFP infection in LMTK and LMTK ρ^0^ cells; cGAS and STING expression were confirmed by western blot (inset). (I) Detection of VSV cDNA in virus-infected cells. N gene-specific primers and probes were used to quantitate VSV cDNAs. GAPDH was used as a control for cellular target DNA. (J) VSV cDNAs are sensitive to DNase I. Cytosolic extracts were incubated with the indicated nucleases, cleaned up, and subjected to qPCR. (K) Detection of GAPDH cDNA. Total cellular DNA was subjected to qPCR with genomic DNA (gDNA)- and splice dependent (cDNA)-specific primer and probe sets.

To more broadly assess cytosolic and hcGAS-bound DNAs, we developed deep sequencing libraries from cytosolic extracts or after immunoprecipitation of WT hcGAS-HA3x. The first one-third of the mitochondrial genome was specifically enriched in cytosolic preps from both uninfected and VSV-GFP-infected MEFs (Figure 5F). Similarly, mtDNA was also highly enriched after immunoprecipitation of hcGAS-HA3x, although there was a bias for the latter three-quarters of the genome (Figure 5G). Importantly, there was no obvious difference in mtDNA pulldown between uninfected and infected cells. Collectively, these data indicate that VSV does not induce cytosolic release of mtDNA to stimulate cGAS activation. Consistent with this, VSV-GFP replicated equally well in LMTK cells and mtDNA-depleted LMTK ρ^0^ cells (41), which express cGAS and STING (Figure 5H). Collectively, these data suggest that mtDNA is dispensable for cGAS-mediated restriction of an RNA virus.

Although VSV is a negative-strand RNA virus that replicates solely via RNA intermediates, it has been reported that VSV-specific cDNAs can arise in infected cells, presumably through reverse transcriptase (RT) activity encoded by endogenous retroelement(s) (42). We therefore investigated whether such viral cDNAs arose during VSV-GFP infections in our laboratory. Indeed, VSV N gene-specific cDNAs were generated in infected cells, although in extremely low abundance, ~1 copy/10^4^ cells (Figure 5I). The cDNA origin of the N gene template was confirmed by nuclease treatment (Figure 5J), by its sensitivity to tenofovir, an RT inhibitor that had no effect on VSV replication (Figure S3A), and by its enhanced expression in cells devoid of TREX1 nuclease (Figure S3B). We also identified virus-specific cDNAs in cells infected with yellow fever virus (YFV), a positive-strand RNA virus (Figure S3C), suggesting that cDNA formation is a general feature of RNA virus infections. Finally, to determine whether cDNA formation was specific to virus-infected cells or to viral transcripts, we examined whether cDNA forms of an abundant housekeeping gene, GAPDH, arose in uninfected cells. Indeed, splice-dependent GAPDH cDNAs were identified in low abundance by qPCR (Figure 5K). Importantly, VSV or retroelement cDNAs were not detected in deep sequencing analyses of whole cytosol or cGAS-HA immunoprecipitations, likely due to their low abundance.

Collectively, our results indicate that cGAS binds mtDNA in both infected and uninfected cells, and that VSV infection does not induce the release of mtDNA into the cytosol or increase cGAS-bound mtDNA. Additionally, viral and cellular mRNA-specific cDNAs can be detected, but are of extremely low abundance, less than one copy per 10^4^ cells. Taken together, these results suggest that steady state levels of cytosolic DNA, rather than virus-induced DNAs, may provide ligands for cGAS-mediated restriction of RNA virus replication.

### cGAS primes smoldering baseline ISG expression

Based on the above results, we hypothesized that cGAS may serve to program baseline levels of innate immune activation rather than strictly in response to RNA virus infection. To address this, we analyzed ISRE-driven luciferase expression in uninfected THP-1 cells devoid of cGAS expression or reconstituted with WT or mutant hcGAS-HA3x (Figure 6A). These experiments suggested that WT hcGAS-HA3x significantly enhances baseline ISG induction compared to the parental cGAS KO line and hcGAS mutants. Further experiments showed that WT hcGAS-HA3x also stimulated greater ISRE-driven luciferase production during infection of THP-1 cells with VSV-GFP and SINV-GFP (Figures 6B, C). It should be noted that VSV-GFP ISG levels were not appreciably different from the control, likely due to the transcriptional repression capability of the M protein (43, 44).

**Figure 6.**
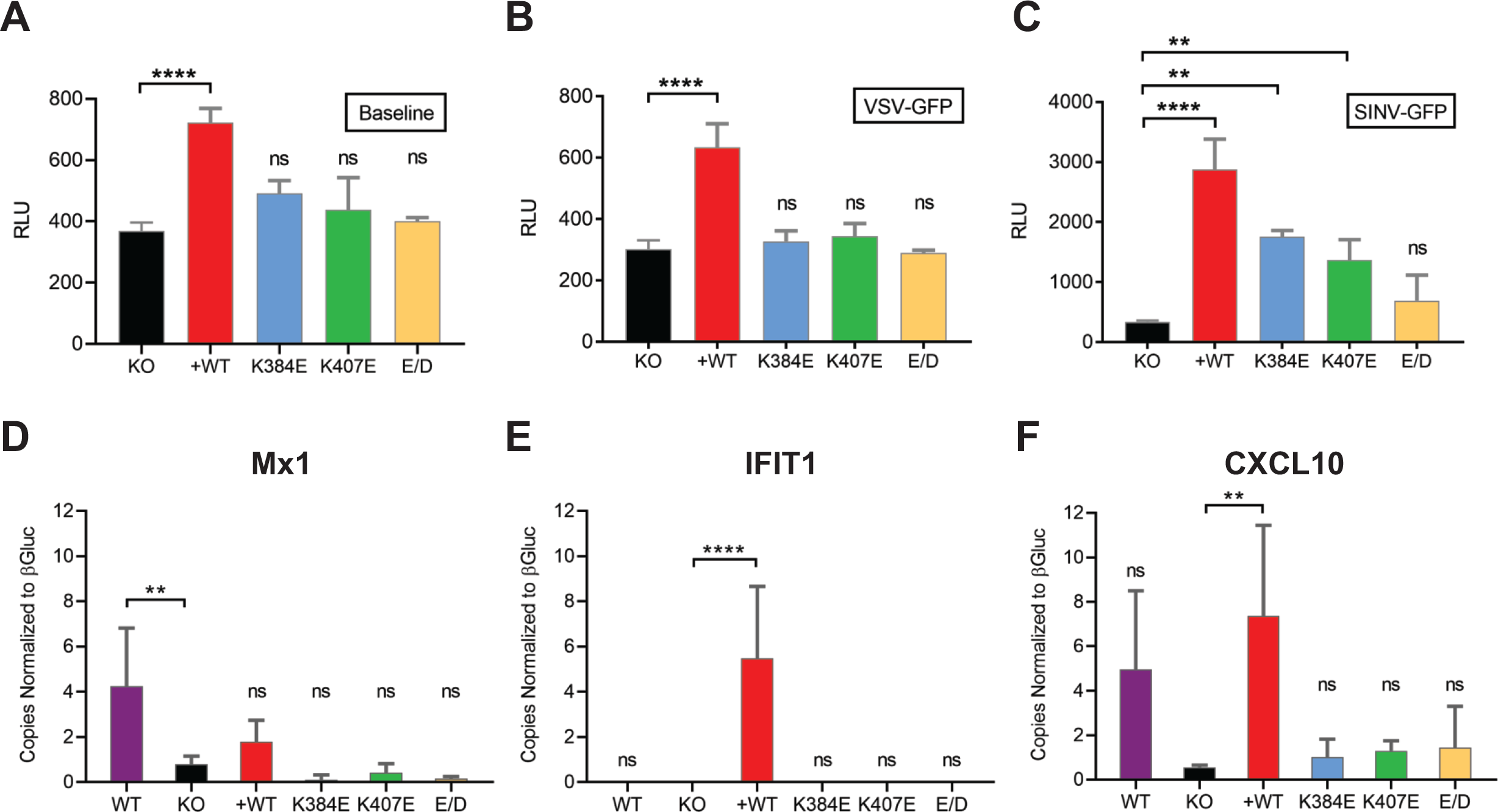
cGAS primes basal ISG expression in steady state cell cultures. ISRE-driven luciferase production in (A) uninfected, (B) VSV-GFP infected, and (C) and SINV-GFP infected THP-1 KO cells lines with or without WT or mutant hcGAS-3xHA expression. RT-qPCR of (D) MX1, (E) IFIT1, and (F) CXCL10 expression in uninfected WT THP-1, THP-1 KO, or THP-1 KO cells expressing WT or mutant hcGAS-HA3x.

To confirm our ISRE-luciferase findings, we used RT-qPCR to quantify ISG transcripts known to be induced by the cGAS-STING DNA sensing pathway (Figures 6D–6G). These results show that cGAS KO significantly reduced basal expression of Mx1 and CXCL10 in uninfected cells, but not of IFIT1, which was not expressed basally. Importantly, cells reconstituted with WT hcGAS-HA3x expressed significantly higher levels of IFIT1 and CXCL10 mRNA, while cells expressing inactive hcGAS-3xHA mutants did not. As these experiments were conducted in cells that slightly overexpressed hcGAS (Figure S2), ISG upregulation likely reflects reinforced, native patterns of expression.

To examine whether cGAS drives basal levels of innate immune activation *in vivo*, we examined ISG expression in vaginal tissue from uninfected WT B6J mice or in mice defective for several innate immunity pathways. As shown in Fig. 7, low basal levels of USP18, Mx1, and Rsad2 expression were observed in B6J mice, but were significantly decreased in IFNAR1^−/−^ mice, demonstrating that basal ISG expression depends on IFNAR signaling. Importantly, cGAS^−/−^ mice had significant decreases in basal Mx1 and Rsad2 expression, similar in degree to reduced basal USP18 and Rsad2 expression observed in IRF3/7^−/−^ mice. In contrast, MAVS had little effect on basal ISG expression.

**Figure 7.**
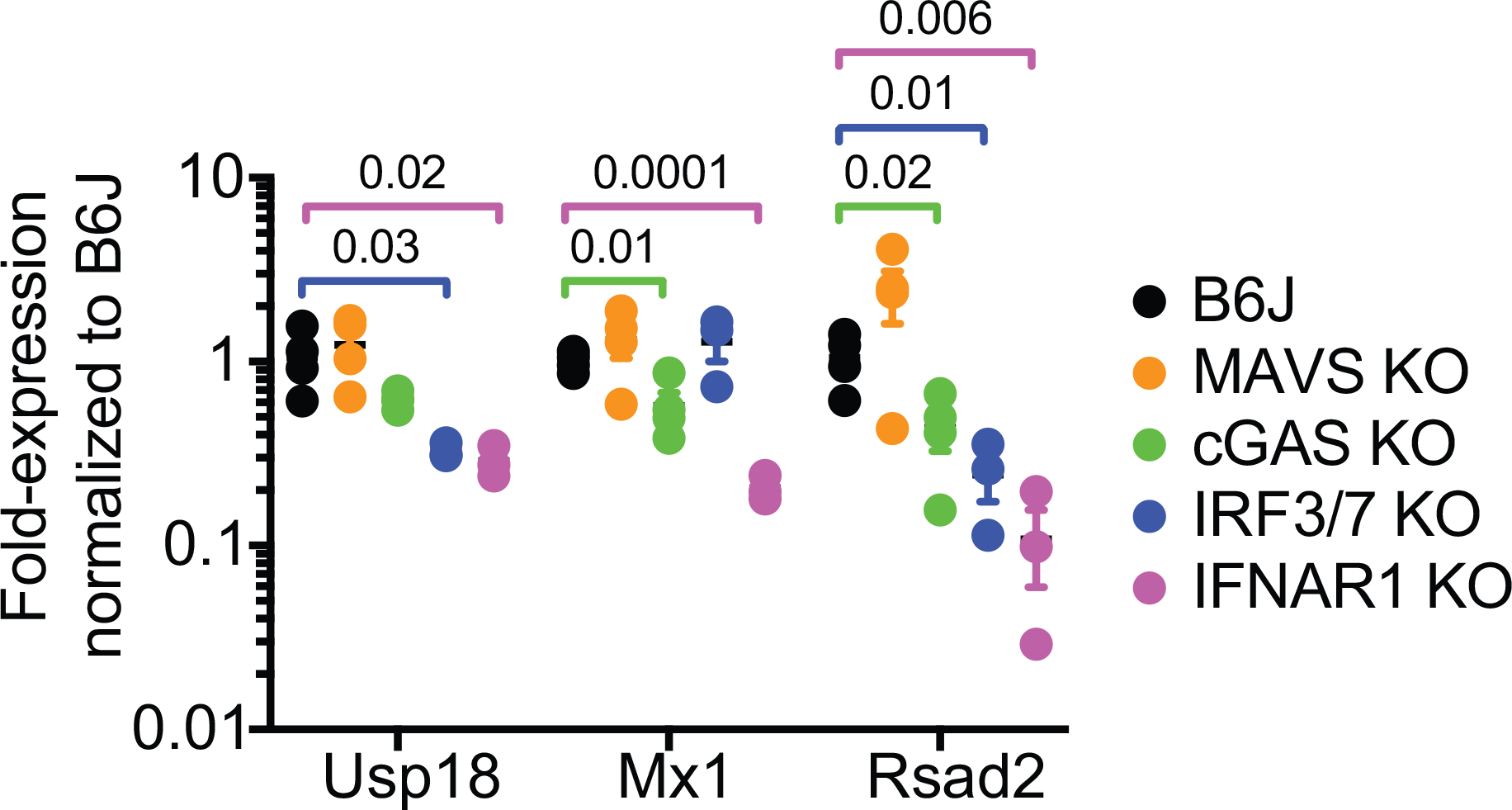
cGAS primes basal ISG expression*in vivo*. Vaginal tissue was collected from uninfected female mice of the indicated genotypes, synchronized in diestrus. Expression levels of USP18, Mx1, and Rsad2 were quantitated by RT-qPCR and normalized to B6J mice.

Altogether, these results suggest that cGAS primes cells to express smoldering levels of ISG expression and that the DNA binding and catalytic activity are integral to this phenomenon.

## Discussion

While the RLR-MAVS and cGAS-STING pathways are important, respectively, for restricting RNA and DNA virus infections, there is considerable crosstalk and redundancy between these two pathways. For instance, mammalian RNA polymerase III can transcribe A-T-rich DNA in the cytosol, producing uncapped RNAs that trigger RIG-I (45, 46). In addition, STING can physically associate with RIG-I and MAVS and may act as a cofactor in RNA sensing (47–49). More recently, STING has been shown to inhibit RNA virus replication, independent of ISG expression, via translational control (20).

Although cGAS was previously reported to restrict RNA viruses (50), it has been widely assumed — though unproven — that this restriction depends on cGAS’s DNA binding and cGAMP synthase activities. Here, we used genetic knockout and transgenic replacement to determine that both DNA binding and cGAMP synthase activities are essential for cGAS-mediated restriction of RNA viruses. One caveat to this approach is that gene knockout can have far-reaching network-level effects on transcription, which are just beginning to be unearthed (51). A second caveat is that reconstituted cGAS was slightly overexpressed in THP-1 cells, which, at least for WT cGAS, can induce ISG expression (50, 52) and may have exaggerated the response. Nevertheless, our results in THP-1 cells were consistent with results obtained from MEFs (Figure 3), which did not overexpress cGAS. Taken together, these data establish that DNA binding and cGAMP synthase activities are required for cGAS-mediated RNA virus restriction.

Despite the essential role of cGAMP synthase activity and demonstrated detection of cGAMP synthesized after DNA transfection, we were unable to detect cGAMP production in response to VSV-GFP infection. Our results are consistent with results recently reported by Franz et al., who were also unsuccessful in detecting cGAMP production in VSV-infected cells (20). While Franz and colleagues concluded that cGAMP is not produced in response to VSV infection, we also considered the possibility that cGAMP levels may be below the limit of detection and/or rapidly degraded. Whereas cGAMP synthesis is readily detected in response to DNA transfection, this may simply reflect the wide dynamic range of cGAS in response to overloading the cytosol with transfected DNA. Moreover, it has been exceedingly difficult to detect cGAMP after virus infections, even for DNA viruses. For instance, Paijo et al. reported that the detection of cGAMP produced in response to cytomegalovirus infection was cell type-dependent, despite active cGAS-STING expression. Where cGAMP was detected, levels were on the order of 5 fmol/10^4^ cells, or ~3×10^5^ molecules/cell, which was slightly above their assay’s limit of detection (53). As our biochemical and biological assays were both less sensitive than that of Paijo et al., we surmise that the synthesis of cGAMP in response to RNA virus infection is below the limit of detection and/or may be rapidly turned over. Alternatively, continuous low-level production of cGAMP in response to endogenous DNA ligands may be more relevant to RNA virus restriction. Clearly, cGAMP assays with improved sensitivity are needed to discern between these possibilities.

Because cGAS DNA binding activity was required for VSV restriction, we examined whether VSV introduces cGAS DNA ligands into the cytosol. Prior work has shown that the cytosolic release of mtDNA activates the cGAS-STING pathway (33–35); moreover, infection with HSV-1, a DNA virus, or dengue virus, an RNA virus, reportedly causes cytosolic release of mtDNA (34, 40). An emerging concept is that mammalian cells may regulate the efflux of mtDNA into the cytosol in response to stress, supported by a role for the Bax/Bak pore in mtDNA release as well as mitochondrial inner membrane release mechanisms via permeabilization and herniation (33–35, 54, 55). In contrast, the levels of cytosolic mtDNA and cGAS-associated mtDNA did not increase during VSV infection. Moreover, cGAS-STING-mediated VSV restriction was intact in ρ^0^ cells, which lack mtDNA, consistent with similar experiments reported by Franz et al. (20). Real-time examination of mitochondrial dynamics may be needed to clarify the role of mtDNA release during RNA virus infections.

Given that cGAS recognizes RNA:DNA hybrids(22), as well as a recent report of VSV cDNAs (42), we also quantitated viral cDNAs produced during VSV infection. We confirmed that rare viral and cellular cDNAs are indeed produced, most likely by an endogenous cellular RT; however, the abundance of any given cDNA was incredibly low, ~1 copy per 10^4^ cells. This was less than the amount of VSV N-gene cDNA previous reported by Shimizu, et al. (42), which we attribute to the enhanced specificity of our hydrolysis probe-based assay vs. SYBR green assays. Nevertheless, the low abundance of VSV cDNAs is inconsistent with a model whereby RNA virus cDNAs play a significant role in stimulating population-wide innate immune responses. These findings, however, do highlight the constant synthesis and turnover of cDNAs within the mammalian cytosol. Consistent with this, deficiencies in the TREX1 nuclease lead to cytosolic accumulation of DNA, including retroelement cDNAs, causing chronic cGAS stimulation and autoimmunity in the form of Aicardi-Goutières syndrome (4, 56, 57).

Given that cGAS may be continuously stimulated by endogenous DNA ligands, and that candidate DNA ligands were unchanged during VSV infection, we examined whether cGAS contributes to a pre-existing baseline of innate immune activation. Indeed, low level cGAS-dependent ISG expression was observed even in the absence of viral infection and was significantly decreased in cGAS KO cells, consistent with prior examples of the cGAS-STING pathway altering ISG baseline expression (2, 50, 58, 59). These results support the hypothesis that cGAS contributes to RNA virus restriction by establishing smoldering, baseline-levels of constitutive innate immune activation. This is an important distinction from other models where cGAS responds to RNA virus-induced release of mtDNA. Additional work will be needed to definitively identify the relevant DNA ligands that activate cGAS; we suggest that pre-existing baseline stimuli should be considered.

While ISG expression served as a convenient and sensitive readout of baseline cGAS-STING activation in our studies, it should be noted that we did not demonstrate that low-level ISG expression directly contributes to RNA virus restriction. On the surface, our results may seem at odds with those of Franz et al., who recently reported that STING restricts RNA viruses, including VSV, in an ISG-independent manner (20). However, we do not exclude the possibility that smoldering cGAS activation may also contribute to ISG-independent mechanisms of virus restriction via STING.

In summary, we propose that cGAS may become activated in response to RNA virus infection, such as by virus-induced mtDNA release, but also contributes to RNA virus restriction via constitutive, low-level innate immune activation, likely via recognition of endogenous DNA ligands (Fig. 8).

**Figure 8.**
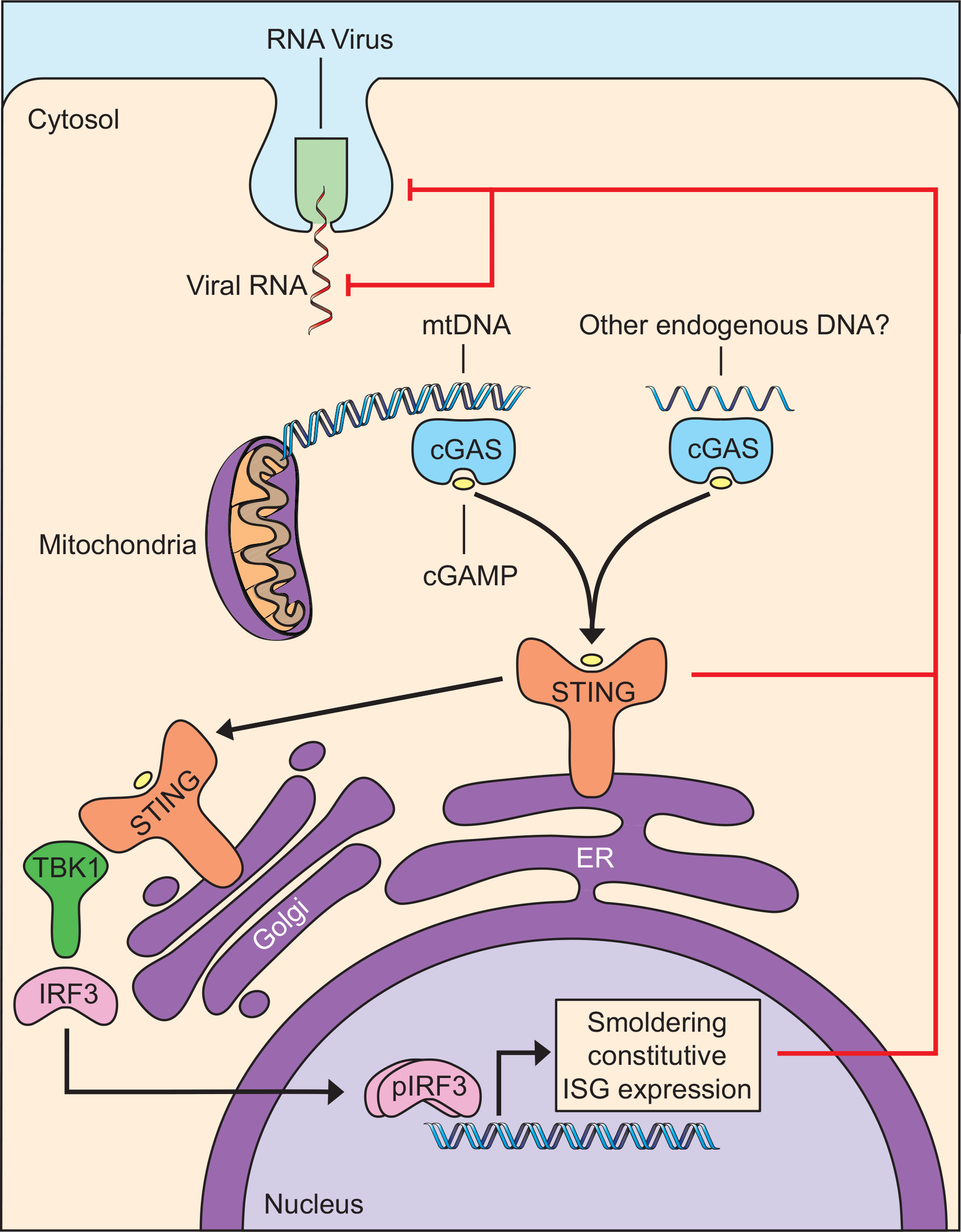
Model of cGAS-mediated restriction of RNA virus infection.

## Materials and Methods

### Animal research

All mice were maintained, bred, and handled in our facility in compliance with federal and institutional policies under protocols approved by the Yale Animal Care and Use Committee. C57BL/6J, B6(C)-Cgas^tm1d(EUCOMM)Hmgu^/J (*cGAS*^−/−^) mice (50) and B6(Cg)-Ifnar1tm1.2Ees/J (*Ifnar1*^−/−^) mice (60) were purchased from Jackson Laboratory. *Irf3*^−/−^ *Irf7*^−/−^ mice (61) were a generous gift by Dr. T. Taniguchi (University of Tokyo) and *Mavs^−/−^* mice (62) were a generous gift by Dr. Z. Chen (University of Texas, Southwestern).

Primary MEFs were isolated from day 14.5 embryos (E14.5) as previously described (63). Vaginal tissue was harvested from six- to twelve-week old female mice synchronized in diestrus via subcutaneous injection with 2 mg Depo-Provera in the neck scruff; mice were sacrificed and vaginal tissue harvested ten days after Depo-Provera treatment.

### Cell cultures

All cells were maintained at low passage by using a seed-lot system and routinely tested for mycoplasma contamination. HEK 293E cells were obtained from Dr. W. Mothes (Yale); SW13 cells were obtained from Dr. C. Rice (Rockefeller University); BHK-21 cells were obtained from Dr. D. Brackney (State of Connecticut Agriculture Research Station); LMTK and LMTK ρ^0^ cells were obtained from Dr. G. Shadel (Yale). HEK293E, SW13, BHK-21 and MEF cells were maintained at 37°C and 5% CO_2_ in complete growth medium (DMEM containing 2 mM L-glutamine [Invitrogen], 10% heat-inactivated fetal calf serum [FCS, Omega Scientific], and 0.1 mM non-essential amino acids [NEAA; Invitrogen]). LMTK and LMTK ρ^0^ cells were maintained in DMEM as above supplemented with 100 μg/mL sodium pyruvate (Invitrogen) and 50 μg/mL uridine (Sigma)

THP-1-Lucia ISG cells (Invivogen) were maintained at 37°C and 5% CO_2_ in RPMI 1640 containing 2 mM L-glutamine, 10% FCS, 0.1 mM NEAA, 10 U/mL penicillin/streptomycin, 100 μg/mL normocin, and 100 μg/mL zeocin.

Pilot experiments showed that THP-1-derived macrophages were more permissive for SINV-GFP than undifferentiated THP-1 monocytes. Differentiation of THP-1 monocytes to macrophages was performed by plating cells at a concentration of 5×10^5^ cells/mL in fresh media and incubating for three days with 100 ng/mL phorbol myristrate acetate (PMA; Invivogen). Adherent monolayers were washed once with DPBS, dissociated with 0.05% trypsin/EDTA, resuspended in fresh media, counted, and seeded for experimentation.

### Viruses

Viruses expressing green fluorescent protein (GFP) were used to facilitate monitoring of virus infections. rVSV-p1-eGFP (VSV-GFP) (64) and rVSV-ΔM51-p5-eGFP (VSVΔM51A-GFP) (65) were kind gifts from Drs. J. Rose and A. van den Pol (Yale), respectively. SINV G100-eGFP (SINV-GFP) (66) was a kind gift from Dr. M. Heise (University of North Carolina, Chapel Hill). VSV and SINV stocks were generated via low multiplicity of infection (MOI = 0.01) passage in BHK-21 cells. Herpes simplex virus 1 KOS-eGFP (HSV-GFP) (67), kindly provided by Dr. P. Desai (Johns Hopkins Medical School), was propagated by low MOI passage (MOI 0.01) in Vero cells, and harvested at 60 hours post-infection. HSV-GFP was prepped by three cycles of freezing (−80°C) and thawing (37°C), clarification (1,500 x g for 15 minutes at 4°C), addition of 10% FCS and 7% dimethyl sulfoxide, aliquoted, and stored at −80°C.

### Plaque assay and fluorescent cell counting

Plaque assays were developed by using semi-solid overlays (DMEM, 10% FCS, 1.6% LE agarose). When plaque formation was evident, cells were fixed with 3% formaldehyde, agarose plugs were removed, and cells stained with 0.1% crystal violet in 20% ethanol. Plaque forming units per mL (pfu/mL) were calculated by counting the number of colonies formed and multiplying this count by the dilution factor.

To prepare GFP-expressing cells for cytometry, cells were trypsinized, washed with DPBS, and fixed in DPBS containing 1% PFA. Fluorescent cells were counted on an Accuri C6 Flow Cytometer (Becton-Dickinson).

### Protein analysis

For western blotting, cells were lysed in RIPA buffer (50 mM Tris pH 8.0, 150 mM NaCl, 1% Triton X-100, 0.5% sodium deoxycholate, 0.1% SDS) containing protease inhibitor cocktail, followed by a 20-minute spin at 16,100 x g and 4°C to remove insoluble material. Protein concentrations were quantified by using a BCA protein assay kit (Thermo Scientific). Equal amounts of protein were separated on 4-12% Bis-Tris Bolt SDS-PAGE gels (Thermo Scientific) and transferred to PVDF membranes by using a Pierce Fast Semi-Dry Blotter. Immunoblotting was performed by 30 minutes of blocking with either 5% milk (American Bio) or SuperBlock (Thermo Scientific) followed by primary antibody and then secondary antibody (2 hours and 1 hour at room temperature, respectively), diluted in the same blocking solution. Blots were developed by using SuperSignal Pico or Femto chemiluminescence substrate kits (Thermo Scientific) and imaged on a GE ImageQuant LAS 4000. Precision Plus protein standards (Bio-Rad) were used to estimate protein molecular weights.

The following primary antibodies were used for western blotting analysis: Rabbit anti-HA (1:1,000, Abcam #ab9110), rabbit anti-pIRF3 (1:1,000, Abcam #76493), rabbit anti-cGAS (1:500, CST #15102s), rabbit anti-TOM40 (1:5,000, Santa Cruz #H-300), rabbit anti-Calreticulin (1:5,000, Abcam #ab2907), rabbit anti-Lamin B1 (1:1,000, Abcam #ab16048), and mouse anti-β-actin (1:10,000, Sigma #A1978). The following secondary antibodies were used for western blotting analysis: Goat anti-rabbit horseradish peroxidase (1:5,000, Jackson ImmunoResearch #111-035-144), and goat anti-mouse horseradish peroxidase (1:5,000, Jackson #115-035-146).

To immunoprecipitate cGAS-DNA complexes, hcGAS-HA3x-expressing cells were fixed in DPBS containing 0.5% paraformaldehyde (5 minutes, room temperature), then quenched with 125 mM glycine. All subsequent steps were performed at 4°C. After two washes with DPBS, cells were lysed for 30 minutes in ice-cold RIPA, followed by a 20-minute spin at 16,100 x g. Clarified lysates were sonicated with four cycles of 10 seconds on and 30 seconds off at 20% amplitude on a Sonifier 450 (Branson Ultrasonics). Samples were spun for 20 minutes at 16,100 x g and supernatants were retained.

To perform immunoprecipitation, lysates were pre-cleared with 2 μg/mL rabbit sera and two incubations with 50 μL protein A-magnetic beads (Pierce). Samples were then rotated overnight with 2 μg HA antibody, and complexes were captured with Protein A-magnetic beads. Washing was performed as follows: 2x with RIPA, 2x with high salt RIPA (500 mM NaCl), 1x with IP-wash buffer (0.5 M LiCl, 1% NP-40, 1% deoxycholate, 100 mM Tris-HCl pH 8.0), and 2x with T_10_E_1_ (10 mM Tris-HCl pH 8.0, 1 mM EDTA). Bound complexes were eluted with 0.1 M glycine-HCl, pH 2.5, samples were neutralized with 1 M Tris-HCl pH 8.0 (0.1 M final), and eluted protein-nucleic acid complexes were then processed for western blotting or deep sequencing (see Nucleic acid purification, below).

### PCR, qPCR, and RT-PCR

Standard PCRs were performed with Phusion DNA polymerase or Taq DNA polymerase (NEB). Unless otherwise noted, cycling was performed for 35 cycles with primers listed in Table 1.

**Table 1.**
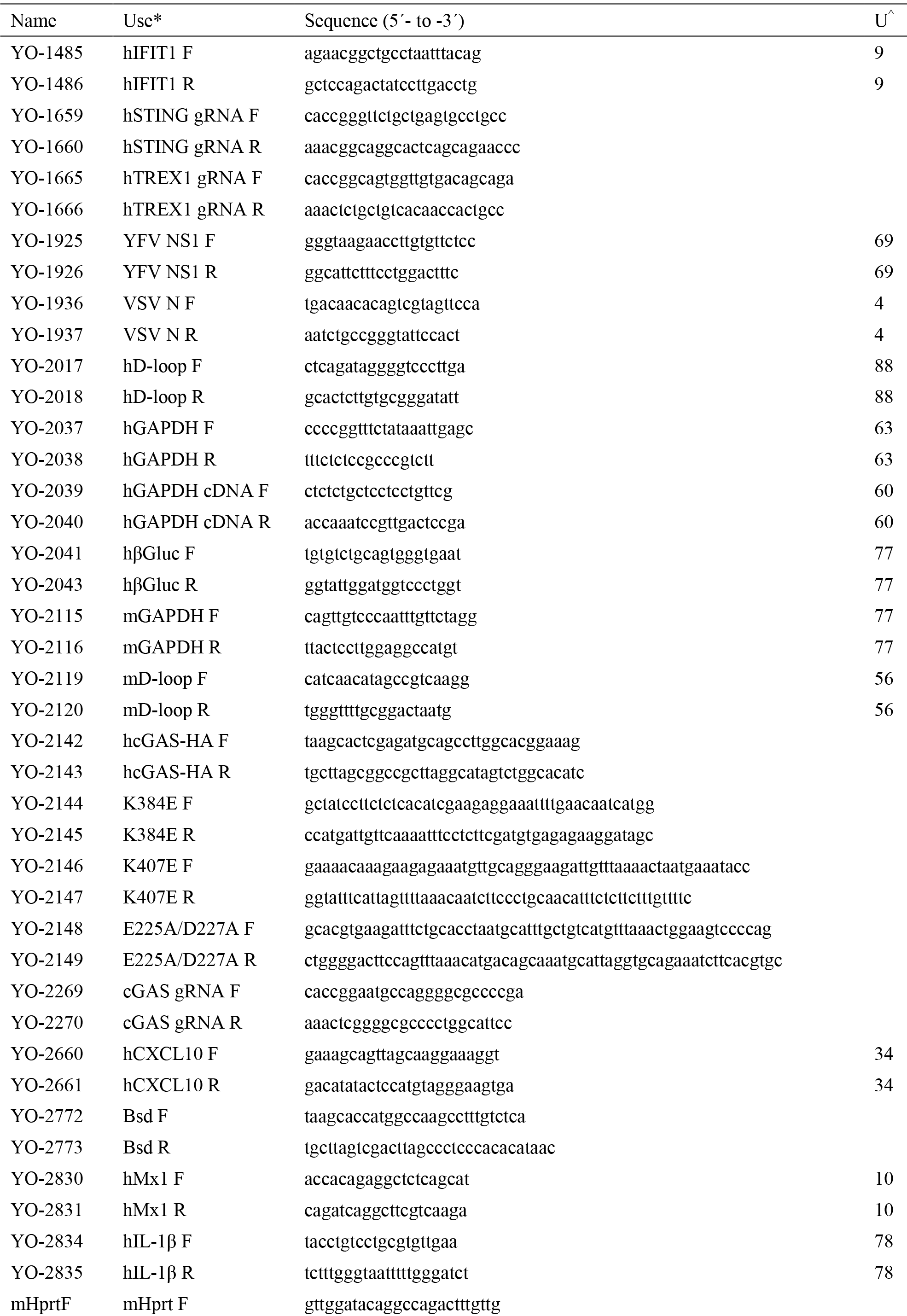
Oligonucleotide primers and probes used in these studies

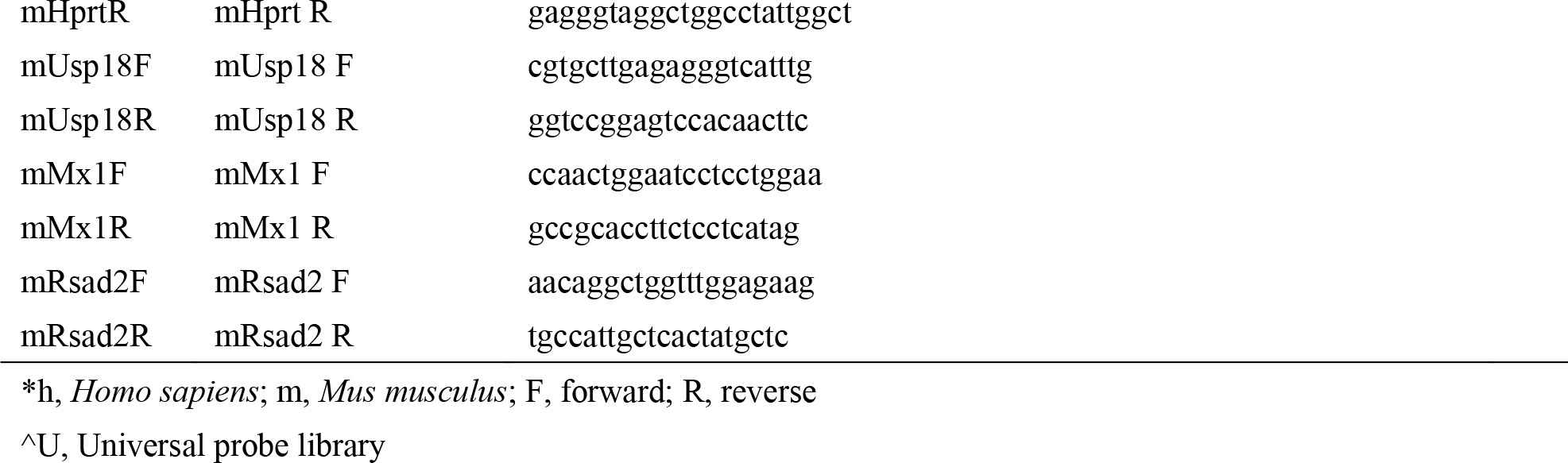

For RT-PCR, RNAs were extracted from cells by using TRIzol Reagent (Life Technologies) or the RNeasy extraction kit (Qiagen). Viral RNA was extracted from cell culture media with the QiAmp Viral RNA Mini Kit (Qiagen). cDNA synthesis was performed by using random hexamer or gene-specific primers with the Transcriptor First Strand cDNA Synthesis Kit (Roche).

For qPCR of cell culture-derived cDNAs, primers were designed by using the ProbeFinder software (Roche) for compatibility with Roche Universal Probe Library (UPL) hydrolysis probes. Assays were performed in a LightCycler 96 or LightCycler 480 (Roche), as per manufacturer’s instructions, with primers and UPL probes listed in Table 1. All reactions were performed in duplicate and quantified by comparison to standard curves created with cloned amplicons diluted (10^2^ – 10^7^ copies) in ddH_2_O supplemented with 50 ng/μL carrier DNA and run in parallel.

For RT-qPCR of mouse tissue-derived mRNAs, SYBR Green qPCR reactions were run in triplicate with gene specific primers (Table 1). The Ct values were averaged, internally normalized against housekeeping gene HPRT, then normalized to B6J control mice by using the ΔΔCt method of comparison. Fold-expression was estimated assuming one doubling per cycle (fold expression = 2^−ΔΔCt^).

### Plasmids, lentiviruses, and retroviruses

pMXs-hcGAS-HA3x-IRES-puro was made by PCR amplifying hcGAS-HA3x from pUNO1-hcGAS-HA3x (Invivogen) with YO-2142 and YO-2143 and cloning into pMXs-mcGAS-HA3x-IRES-puro (gift of Dr. G. Shadel, Yale University) via common XhoI and NotI sites. To facilitate reconstitution of puromycin-resistant cGAS knockout (KO) cells with hcGAS, the Pac gene in pMXs-hcGAS-HA3x-IRES-puro was replaced with the Bsd gene, amplified from pMICU-APEX2 (68) (Addgene plasmid # 79057). In addition, the 518-bp NotI– BlpI fragment of hcGAS was recoded (BlueHeron) to avoid editing of the reconstituted hcGAS. For transient expression of hcGAS in primary MEFs, hcGAS-3xHA was cloned into pLenti-puro (69) (Addgene plasmid # 39481).

Site-directed mutagenesis of hcGAS was performed by using appropriate primers (Table 1) and PfuTurbo (Agilent Technologies), as previously described (70). Mutants were sequenced and subcloned back into the pMXs-IRES-puro vector with XhoI and NotI. Gene knockout was performed in cell culture by using Cas9 to induce non-homologous end-joining repair. Briefly, gRNAs-specific oligos (Table 1) were chosen from published datasets (71) or designed with gRNA Designer (72) and cloned into pLentiCRISPR (73) (Addgene plasmid # 51760).

### Lentiviruses and retroviruses

Lentiviruses and retroviruses were packaged in HEK 293E cells by co-transfection with appropriate HIV- or MLV-Gag/Pol and VSV G packaging constructs. Forty-eight hours post-transfection, packaged vector stocks were clarified (16,100 x g), passed through a 0.45 μm filter, and supplemented with 8 μg/mL polybrene (Sigma) and 20 mM HEPES (Life Technologies). Target cells were transduced by spinoculation, selected with 3 μg/mL puromycin, and screened for expression or knockout via genomic PCR and sequencing and/or western blotting. To develop clonal cultures, adherent cells were isolated by using sterile 8 mm Pyrex cloning cylinders and expanded. Clonal phenotypes were screened via western blotting or qPCR.

### Luciferase assays

Lucia luciferase-containing samples were clarified by centrifugation (16,100 × g for 5 minutes) and mixed with ¼-volume of 5x Renilla lysis buffer (Promega) to destroy virus infectivity. Lucia activity in 20 μL samples was measured on a Centro LB 960 plate reader (Berthold) by integrating over 10 seconds.

### cGAMP extraction and assays

Cells were infected for five hours with VSV-GFP or SINV-GFP (MOI 3–10), transfected for five hours with salmon sperm DNA (2 μg/mL final concentration) and Transit LT-1 (Mirus), or left untreated. cGAMP was extracted based on established methods (21). Briefly, cells were dissociated with trypsin, washed with DPBS, gently pelleted and resuspended at a concentration of 1×10^7^ cells/mL in ice-cold cGAMP homogenization buffer (10 mM Tris-HCl, pH 7.4, 10 mM KCl, 1.5 mM MgCl_2_). Cells were lysed via nitrogen cavitation in a cell disruptor (Parr Instrument Company). Lysates were clarified at 1,000 x g for 5 minutes, then 16,100 x g for 10 minutes, retaining the supernatants. Resulting supernatants were digested for 1 hour with benzonase (0.5 U/μL; Fisher Scientific) at 37°C, 1 hour with proteinase K (0.5 U/μL; Invitrogen) at 55°C, heat-inactivated at 95°C for 10 min, and spun for 5 minutes at 16,100 x g, retaining the final supernatant (S1).

To detect cGAMP bioactivity, 2 μL of S1 sample, synthetic cGAMP (positive controls), or DPBS (negative controls) were incubated with 10^6^ THP-1 cells, 2 mM ATP, 1.5 ng/μL SLO (a kind gift from Dr. N. Andrews, University of Maryland) and media in 8 μL (total volume). After 1.5 hours at 30°C, reactions were lysed with an equal volume of RIPA buffer and processed for phosphorylated IRF3 western blot, as above.

To detect cGAMP by liquid chromatography and mass spectrophotometry, trypsinized cells were washed once with DPBS, pelleted at 1,000 x g and then frozen at -20°C. To extract cGAMP, 5×10^6^ cells/mL were resuspended three times in extraction buffer (40% acetonitrile, 40% methanol, 20% ddH_2_O) for 20 minutes, spinning after each extraction at 16,100 x g and keeping the supernatant. Supernatants were pooled, dried overnight in a GeneVac HT-8 (SP Scientific), and resuspended in 100 μL ddH_2_O per 5×10^6^ cells. Samples were filtered with a 0.2 μm PTFE syringe filter (VWR) prior to loading into a Luna Omega C18 UHPLC column (Phenomenex) on an iFunnel 6550 Q-TOF / MS (Agilent). Samples were run in negative mode with the following parameters: Buffer A = 0.1% formic acid; Buffer B = acetonitrile, 0.1% formic acid; gradient cycles: 0 – 4% B over 10 minutes, 4% B – 100% B over 5 minutes, 5 minutes wash with 100% B; UV detection at 260 nm, m/z scans from 150-1,000. cGAMP was observed between 4–7.5 minutes in extracted ion chromatographs at an observed mass of 673.085 m/z; this was confirmed to be cGAMP by MS/MS ion fragmentation patterns.

### Preparation of cytosolic nucleic acid extracts

Cells were trypsinized and resuspended in an equal volume of fresh media, then spun at 1,000 x g for 5 minutes at room temperature. After washing once with DPBS, cells were resuspended in cytosolic extraction buffer (50 mM HEPES pH 7.4, 150 mM NaCl, 25 μg/mL digitonin) and incubated for 10 minutes at 4°C with rotation. A succession of 4°C spins was performed, retaining the supernatant for each step: 3x 1,000 x g for 3 minutes, 1x 16,100 x g for 10 minutes, 1x 100,000 x g for 1 hour on a 0.34 M sucrose cushion (SW41 Ti rotor, Beckman). The final supernatant was then processed for western blotting, above, and DNA purification, below.

### DNA purification and phi29 amplification

DNA was isolated from total cytosol by treating samples with RNase A and RNase T1 (Ambion) for 1 hour at 37°C, digesting with Proteinase K for 1 hour at 55°C, and heat inactivating at 95°C for 15 minutes. DNAs were then purified with the QiaQuick purification kit (Qiagen). To isolate DNA from immunoprecipitates, crosslinks were reversed by adding 5 M NaCl (0.3 M final) and shaking overnight at 65°C, then digesting RNA and protein, as above. For isothermal DNA amplification, 1–40 ng of DNA was annealed to exo-resistant random hexamer primers (Molecular Cloning Laboratories) and amplified overnight at 30°C with phi29 DNA polymerase (NEB), followed by a 65°C inactivation step. DNA was extracted with the QiaQuick purification kit.

### Sequencing library preparation, sequencing, and analysis

Phi29-amplified samples were sonicated, as above, to achieve DNA fragments of 200- to 500-bp. DNAs were end-repaired with T4 DNA polymerase (NEB), T4 polynucleotide kinase (NEB), and Klenow DNA polymerase (NEB) at 20°C for 30 minutes, then purified via QiaQuick. A-tailing was performed with Klenow fragment (3’-5’ exo[-], NEB) before ligation of TruSeq Adapters (Illumina) and amplification with Phusion DNA Polymerase (NEB) and Illumina TruSeq primer cocktails. Size-selected libraries (350- to 450-bp) were excised from LMP agarose, purified with a Gel Extraction Kit (Qiagen), re-amplified by PCR for 18 cycles, then purified. Triplicate libraries were sequenced on a HiSeq4000 (Illumina) by the Yale Center for Genome Analysis with a 150-bp paired-end protocol, 100 million reads/sample.

### Software and Statistics

Unless otherwise noted, statistical significance was estimated by using the Student’s t-test with Holm-Sidak correction for multiple comparisons. Standard p-value indicators were used throughout the manuscript: * indicates p<0.05; ** indicates p<0.01; *** indicates p<0.001; and **** indicates p<0.0001. Data were graphed by using Graphpad Prism software (version 7.0a). Pixel densities were analyzed in ImageJ2 (74) and images were prepared for presentation with Photoshop and Illustrator CS4 (Adobe). Next-generation sequencing results were mapped to the mm10 mouse genome by using the Burrows-Wheeler aligner (BWA) (75) and TopHat (76) to look for raw and gapped alignment, respectively. Alignments were assessed for content of genomic DNA and mtDNA with the integrated genome browser software (77).

## Acknowledgments

We thank Drs. H. Ramanathan, D. DiMaio, and P. Cresswell for constructive feedback; Ms. H. Dong for technical help in isolating primary MEFs; Drs. G. Shadel for SV40 T-immortalized MEFs; Drs. D. Schatz, G. Teng, and S. Mehta for technical help in sequence library preparation and analysis; Drs. J. Rose and A. van den Pol for VSV-GFP; Dr. M. Heise for SINV-GFP; Dr. P. Desai for HSV-1; and Dr. N. Andrews for SLO. This research was funded by 5R01AI087925 (to BDL) and 1R01AI131518 (to BDL), the Yale Interdisciplinary Immunology Training Program (NIH T32AI07019, to Dr. D. Schatz (Yale) in support of MTP), the Yale Gruber Science Fellowship (to MTP), and the National Science Foundation Graduate Research Fellowship (DGE1122492, to MTP).

---

**Figure S1. hcGAS recoding, gRNA binding site, and residues targeted for mutations.** Site-directed mutagenesis was utilized to generate three mutants (brown). The K384E and K407E mutations disrupt the DNA-binding ability of the cGAS, while the E225A/D227A mutation ablates cGAMP catalytic activity. The 5′ 518-bp of hcGAS were codon optimized (red) to improve expression and to generate a sequence resistant to CRISPR/Cas9 targeting. The gRNA binding site (blue) was modified at 8 residues.

**Figure S2. hcGAS KO in THP-1 cells and reconstitution with hcGAS-HA3x**. Western blotting of (A) a dilution curve of THP-1 WT lysate inputs and (B) lysates of THP-1 WT and hcGAS KO cells reconstituted with hcGAS. (C) Standard curve generated from a dilution series of the lysate used in (A). (D) Calculation of relative hcGAS expression levels in (B) as compared to the THP-1 WT sample by using the standard curve in (C).

**Fig. S3. Detection of viral cDNAs.** (A) Time-course of VSV growth and cDNA formation in HEK-293T cells treated with tenofovir and infected with VSV-GFP (MOI 3). Tenofovir (1 μM), which was added one hour prior to infection and maintained throughout the time-course, had no effect on VSV replication. (B) Accumulation of VSV N-gene cDNA in WT HEK 293 cells or HEK 293E cells ablated for TREX1 by CRISPR/Cas9. This experiment was repeated once with similar results. (C) Detection of YFV cDNA. Shown here is the standard curve used to estimate absolute DNA copies via qPCR (left) and the quantity of viral cDNA detected in BHK-21 or Huh-7.5 cells infected with YFV-17D (right). This experiment was repeated twice with similar results.

